# Complex responses to inflammatory oxidants by the probiotic bacterium *Lactobacillus reuteri*

**DOI:** 10.1101/605881

**Authors:** Poulami Basu Thakur, Abagail R. Long, Benjamin J. Nelson, Ranjit Kumar, Alexander F. Rosenberg, Michael J. Gray

## Abstract

Inflammatory diseases of the gut are associated with increased intestinal oxygen concentrations and high levels of inflammatory oxidants, including hydrogen peroxide (H_2_O_2_) and hypochlorous acid (HOCl), which are antimicrobial compounds produced by the innate immune system. This contributes to dysbiotic changes in the gut microbiome, including increased populations of pro-inflammatory enterobacteria (*Escherichia coli* and related species) and decreased levels of health-associated anaerobic Firmicutes and Bacteroidetes. The pathways for H_2_O_2_ and HOCl resistance in *E. coli* have been well-studied, but little is known about how commensal and probiotic bacteria respond to inflammatory oxidants. In this work, we have characterized the transcriptomic response of the anti-inflammatory, gut-colonizing Gram-positive probiotic *Lactobacillus reuteri* to both H_2_O_2_ and HOCl. *L. reuteri* mounts distinct responses to each of these stressors, and both gene expression and survival were strongly affected by the presence or absence of oxygen. Oxidative stress response in *L. reuteri* required several factors not found in enterobacteria, including the small heat shock protein Lo18, polyphosphate kinase 2, and RsiR, an *L. reuteri*-specific regulator of anti-inflammatory mechanisms. These results raise the intriguing possibility of developing treatments for inflammatory gut diseases that could sensitize pro-inflammatory enterobacteria to killing by the immune system while sparing anti-inflammatory, health-associated species.

**IMPORTANCE:** It is becoming increasingly clear that effective treatment of inflammatory gut diseases will require modulation of the gut microbiota. Preventing pro-inflammatory bacteria from blooming while also preserving anti-inflammatory and commensal species is a considerable challenge, but our results suggest that it may be possible to take advantage of differences in the way different species of gut bacteria resist inflammatory oxidants to accomplish this goal.

## INTRODUCTION

Inflammatory diseases of the gut (*e.g.* inflammatory bowel disease [IBD], Crohn’s disease, irritable bowel syndrome [IBS], *etc.*) are a rapidly growing health concern for which few effective treatment options are available (1–3). It has become increasingly clear that the bacterial populations inhabiting the gut play a key role in causing and perpetuating gut inflammation, with an emerging consensus that blooms of facultatively anaerobic enterobacteria (*e.g. Escherichia coli*) take advantage of changes in the nutritional and redox environment of the inflamed gut to outcompete the obligate anaerobes associated with a healthy gut flora (4–8). The redox changes in the inflamed gut include not only increases in oxygen levels (6, 7), but also the production of reactive oxygen, nitrogen, and chlorine species (ROS, RNS, RCS), which are antimicrobial oxidants that can shift the population structure of the microbiome and are major contributors to host tissue damage (9–12). Treatments that interfere with the ability of enterobacteria to thrive in the inflamed gut reduce both the changes in the microbiome and the symptoms of disease (1, 13), indicating that manipulating gut bacteria is an important element in controlling these diseases.

Probiotics are live microorganisms which, when consumed in sufficient quantities, have a measurable health benefit (14), and a variety of different probiotic bacteria have been shown to have anti-inflammatory effects in the gut (15, 16). The most commonly used probiotics are lactic acid bacteria of the genus *Lactobacillus* (17), which are able to both modulate the host immune system and outcompete enterobacterial pathogens (15), and some strains of which have been shown to improve outcomes for inflammatory bowel diseases in both humans and animal models (18–20). The effectiveness of probiotics for treating inflammation in the gut, however, may be limited by their ability to survive attack by the overactive host immune system, including the oxidative damage caused by ROS and RCS. While the general stress response physiology of lactic acid bacteria has been relatively well characterized (21), bacterial responses to oxidative stress are best understood in *E. coli* and related inflammation-enriched enterobacteria (22–25). This is especially true of RCS, including hypochlorous acid (HOCl) and reactive chloramines, which are extremely potent antimicrobial compounds produced by the neutrophil enzyme myeloperoxidase (22, 26–28). Relatively little is known about how health-associated probiotic and commensal bacteria sense and respond to inflammatory oxidants (21, 29–31).

*Lactobacillus reuteri* is a well-established model probiotic bacterium that is able to stably colonize the mammalian intestine (32, 33), where it combats inflammation and enteric infections by several different mechanisms, including anti-inflammatory histamine synthesis (34–37), modulation of immune cell functions (33, 38–40), and production of antimicrobial compounds (*e.g.* reuterin, reutericyclin)(37, 41, 42). While the genome-wide stress responses of *L. reuteri* to low pH (43) and bile salts (44, 45) have been characterized, little is known about how this organism responds to ROS, and nothing is known about how *L. reuteri* or any other lactic acid bacterium senses or responds to RCS. The *L. reuteri* genome encodes neither catalase nor superoxide dismutase (49). The oxidative stress repair enzyme methionine sulfoxide reductase (46) is induced by and required for gut colonization by *L. reuteri* (47, 48), indicating that resistance to oxidative damage is important *in vivo*. A cysteine-dependent pathway contributing to H_2_O_2_ and O_2_ tolerance has been identified (50), but did not appear to play a role in the ability of *L. reuteri* to prevent colitis (51).

In this work, we have now taken a transcriptomic approach to characterize genome-wide H_2_O_2_- and HOCl-dependent gene regulation in *L. reuteri* and to identify genes involved in resistance to killing by these stressors (52), with the goal of finding genes and pathways distinct from those found in the enterobacteria. Our results show that, despite not containing close homologs of any of the known RCS-specific transcription factors (22, 53–56), *L. reuteri* is able to mount clearly different stress responses to H_2_O_2_ and HOCl stress, and that the presence of O_2_ has dramatic effects on both gene regulation and survival in response to these stresses. We also identified roles for several genes in surviving H_2_O_2_- and HOCl-mediated killing, including those encoding methionine sulfoxide reductase (46), polyphosphate kinase 2 (57, 58), and the lactic acid bacteria-specific small heat shock protein Lo18 (59–61), as well as an unexpected role in surviving H_2_O_2_ stress for RsiR, previously characterized as an *L. reuteri*-specific regulator of histamine synthesis (35). Ultimately, we hope these results will help lay the groundwork for the development of targeted treatments for inflammatory gut diseases that could either preferentially sensitize disease-associated enterobacteria to killing by the immune system or preferentially protect health-promoting probiotic and commensal species, thereby stabilizing the healthy microbiome against inflammatory stress.

## RESULTS AND DISCUSSION

### Growth of *L. reuteri* is inhibited by inflammatory oxidants

To begin characterizing the response of *L. reuteri* to inflammatory oxidants, we treated anaerobically growing cultures with different concentrations of H_2_O_2_ (Fig. 1A) and HOCl (Fig. 1B). *L. reuteri* growth was more sensitive to H_2_O_2_ than HOCl, with nearly complete inhibition by 0.96 mM H_2_O_2_ or 5 mM HOCl. The sensitivity of *L. reuteri* to H_2_O_2_ was very similar to that reported for *L. acidophilus*, another catalase-negative probiotic (31), but, as expected, *L. reuteri* was considerably more sensitive to H_2_O_2_ than the pseudocatalase-expressing species *L. plantarum* (62, 63). Since we were interested in characterizing gene regulation during a successful, productive response to sublethal stress, we selected concentrations of 0.12 mM H_2_O_2_ and 1.25 mM HOCl for further analysis. These concentrations resulted in very similar slight reductions in growth rate after stress treatment, followed by complete recovery (Fig. 1A and B, red circles). These concentrations of oxidants had no significant effect on cellular NAD^+^ / NADH ratios (Fig. 1C, 1D, and S1), indicating that the toxic effects of sublethal H_2_O_2_ and HOCl stress did not include major disruptions to the redox state of the bacterial cells.

**FIG 1.**
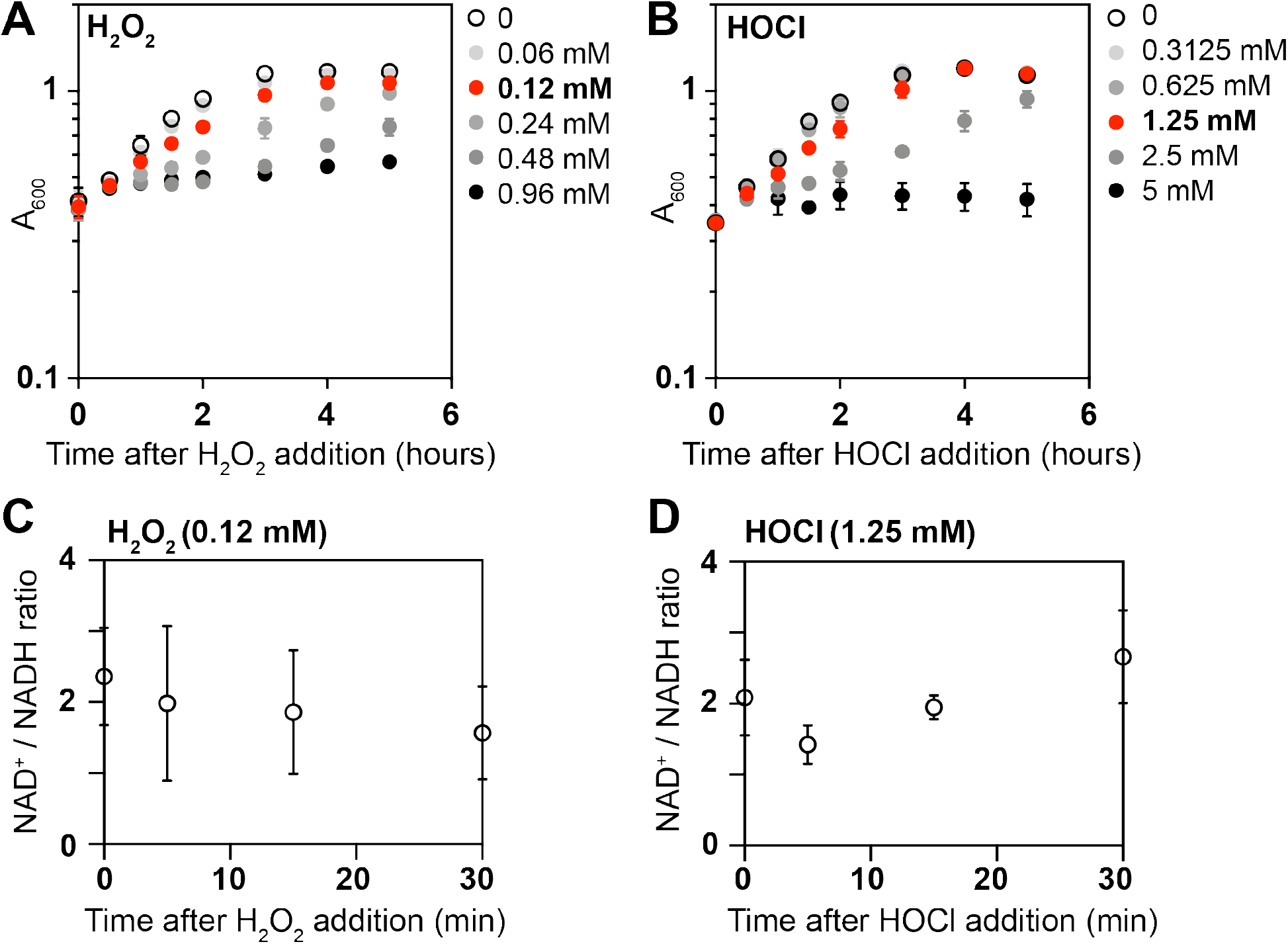
Growth of *L. reuteri* is inhibited by inflammatory oxidants. *L. reuteri* ATCC PTA 6475 was grown anaerobically at 37°C to A_600_=0.3–0.4 in MEI-C and then treated with the indicated concentrations of H_2_O_2_ (*A, C*) or HOCl (*B, D*) (n=3, ±SD). A_600_ (*A, B*) or NAD^+^ / NADH ratios were measured at the indicated times. Red symbols indicate the stress treatments used for subsequent transcriptomic analyses.

### Transcriptomic analysis of H_2_O_2_ and HOCl response by *L. reuteri*

We next treated anaerobically growing *L. reuteri* with sublethal doses of H_2_O_2_ and HOCl and used RNA sequencing to characterize the transcriptomes of stressed cells before and 5, 15, and 30 minutes after stress treatment (Fig. 2, Table S1). A very large fraction of the *L. reuteri* genome was up- or down-regulated (> 2-fold, p < 0.01) by sublethal oxidative stress, and there were clear differences in the responses to H_2_O_2_ and HOCl, consistent with previous reports that bacterial responses to these oxidants are different (22–24). As seen in Figure 2, the response to H_2_O_2_ involved roughly equal numbers of up- and downregulated genes, with a substantial increase in the number of genes with significant changes in expression over the 30 minute course of stress treatment. In contrast, HOCl treatment caused many more genes to be upregulated than downregulated, and there was not as noticeable an increase in the number of genes with significant changes in gene expression over time, consistent with the very fast reaction rate of HOCl with biological molecules (22, 27, 64). The difference between the H_2_O_2_ and HOCl stress responses were also reflected in principal components analysis of the transcriptomic data (Fig. S2A), which clearly separated the H_2_O_2_ and HOCl treated samples. The untreated samples from the two experiments did not cluster as closely together as we expected, since these samples were ostensibly identical. To determine whether this reflected batch effects or inherent variation in expression levels for particular genes, we selected representative genes that had similar (*sigH, moeB, pcl1*) or different (*pstS, copR*) levels of expression in the untreated samples from the two RNA-seq data sets (Table S1) and used RT-PCR to measure their expression in independently prepared unstressed *L. reuteri* cultures, and found that the amount of variation in expression was similar for all five genes (Fig. S2B).

**FIG 2.**
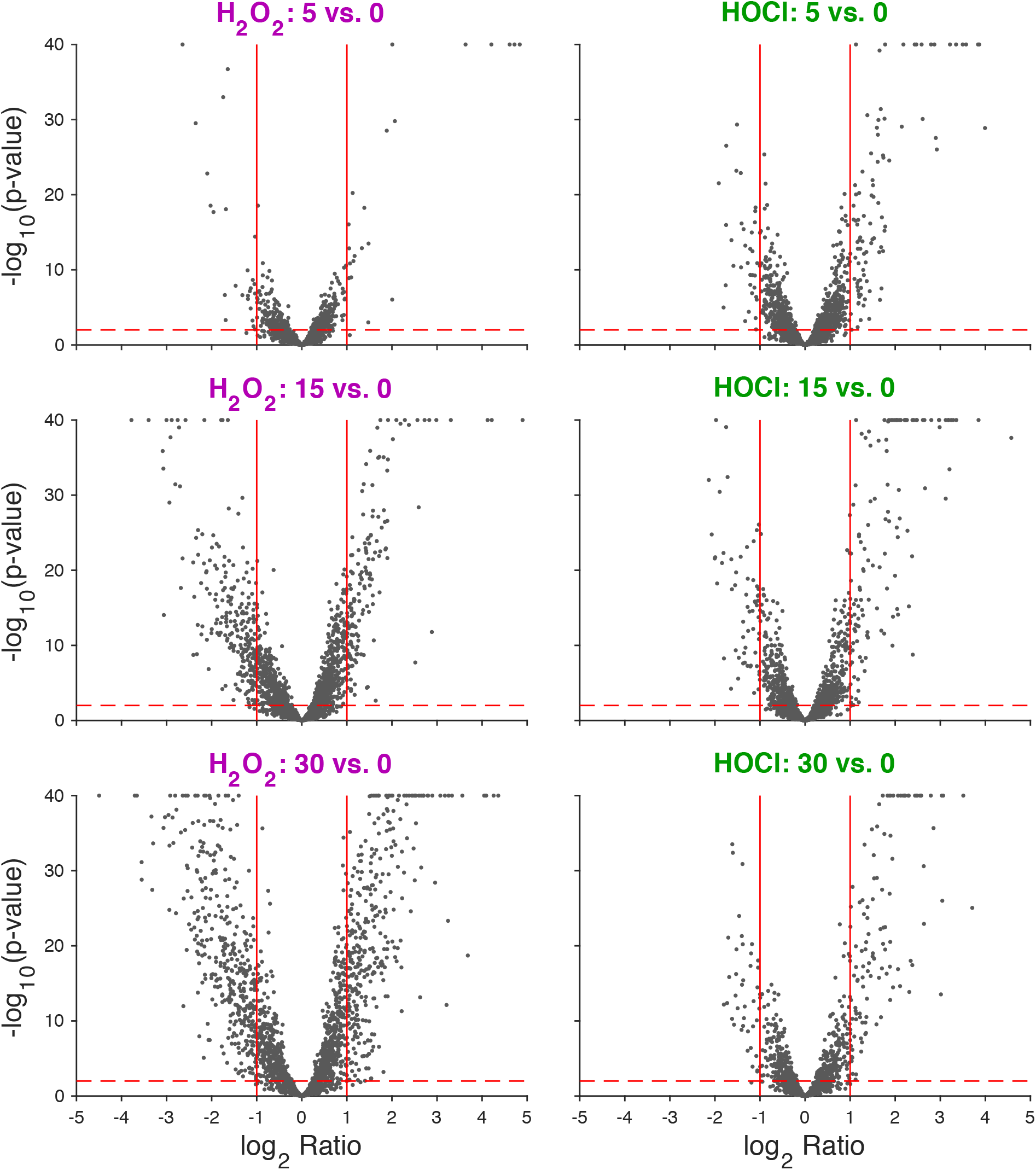
Volcano plot of log_2_-fold changes in gene expression in anaerobically-grown *L. reuteri* ATCC PTA 6475 treated with 0.12 mM H_2_O_2_ or 1.25 mM HOCl 5, 15, and 30 minutes after stress treatment. −log_10_ p-value is plotted against log_2_ gene expression ratio for H_2_O_2_-treated (left column) and HOCl-treated (right column) cultures for three time-points relative to their baseline (rows). Vertical red lines indicate +/- 2-fold changes. Horizontal red dashed line corresponds to p = 0.01.

To further characterize the differences and overlaps between the H_2_O_2_ and HOCl stress responses of *L. reuteri*, we plotted changes in gene expression under each tested condition against each other condition (Fig. 3, Table S2). This showed that, while many of the genes significantly upregulated by H_2_O_2_ were also upregulated by HOCl, there were substantial numbers of genes that were upregulated by one stress treatment and downregulated by the other and *vice versa*, indicating that *L. reuteri* has a sophisticated ability to distinguish between H_2_O_2_ and HOCl and differentially control transcription. Clustering analysis of genome-wide expression patterns (Fig. 4) reinforced this result, and we were able to identify groups of genes whose expression was controlled in very similar ways by the different oxidants (*e.g.* Fig. 4B and G) as well as groups of genes with very different expression patterns in response to H_2_O_2_ and HOCl (*e.g.* Fig. 4C, D, and H), including, for example, *cgl* and *cyuABC*, which encode a previously characterized cysteine-dependent redox stress response pathway (50). This is consistent with results in *E. coli* and *Bacillus subtilis*, in which H_2_O_2_ and HOCl stress responses partially overlap, but have substantial oxidant-specific components (22, 27, 56, 64, 65).

**FIG 3.**
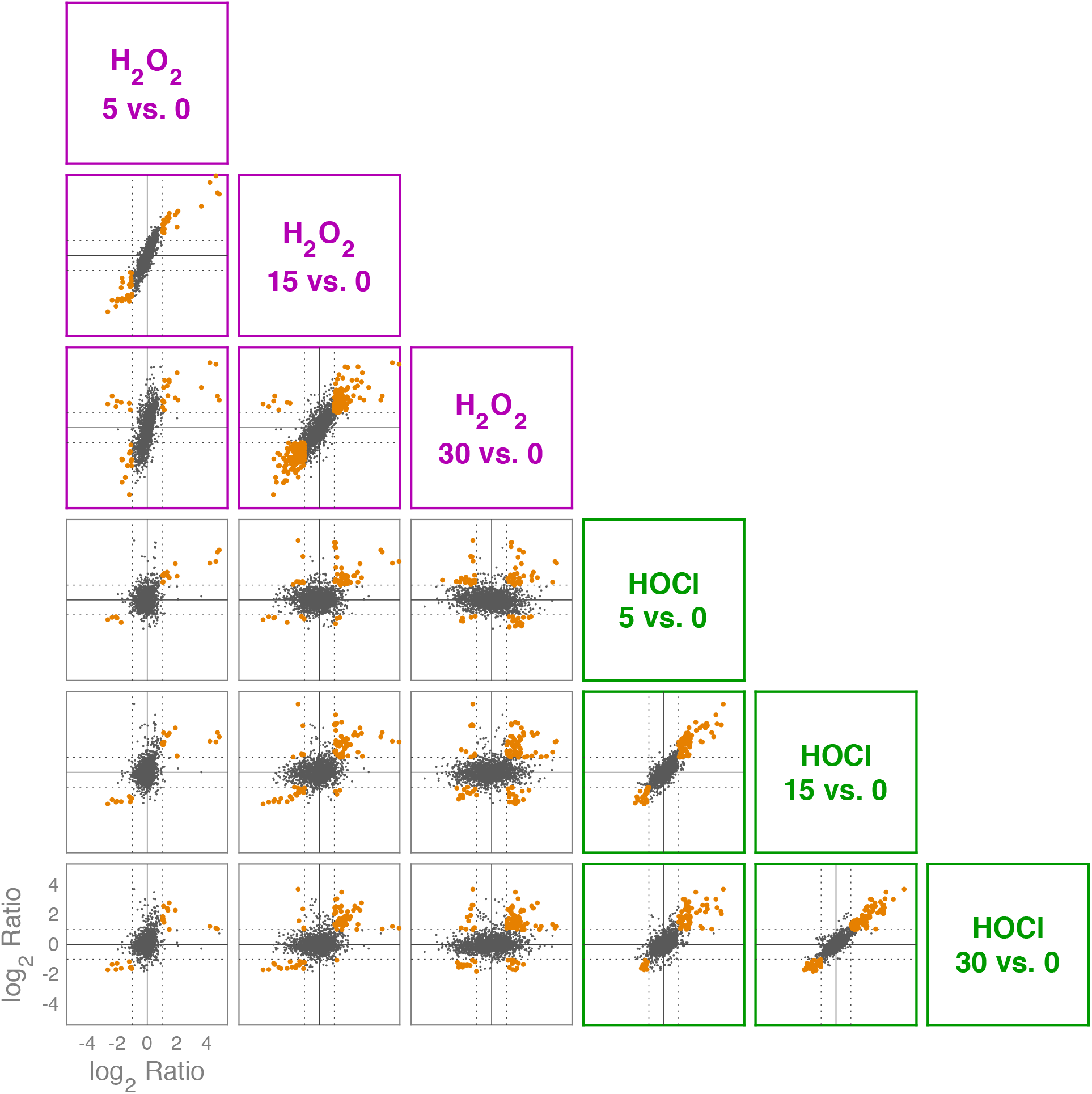
Log_2_-fold changes in gene expression in anaerobically-grown *L. reuteri* ATCC PTA 6475 treated with 0.12 mM H_2_O_2_ (purple) or 1.25 mM HOCl (green) 5, 15, and 30 minutes after stress treatment. 15 pairwise comparisons of log_2_ ratios of six differential expression experiments. Each dot represents a gene and its position reflects the log_2_ ratio in each of two differential expression results. Horizontal lines represent +/- 2-fold changes. Orange points are genes with 2-fold changes in both differential expression data sets (not necessarily in the same directions). See Table S2 for individual genes in each expression category.

**FIG 4.**
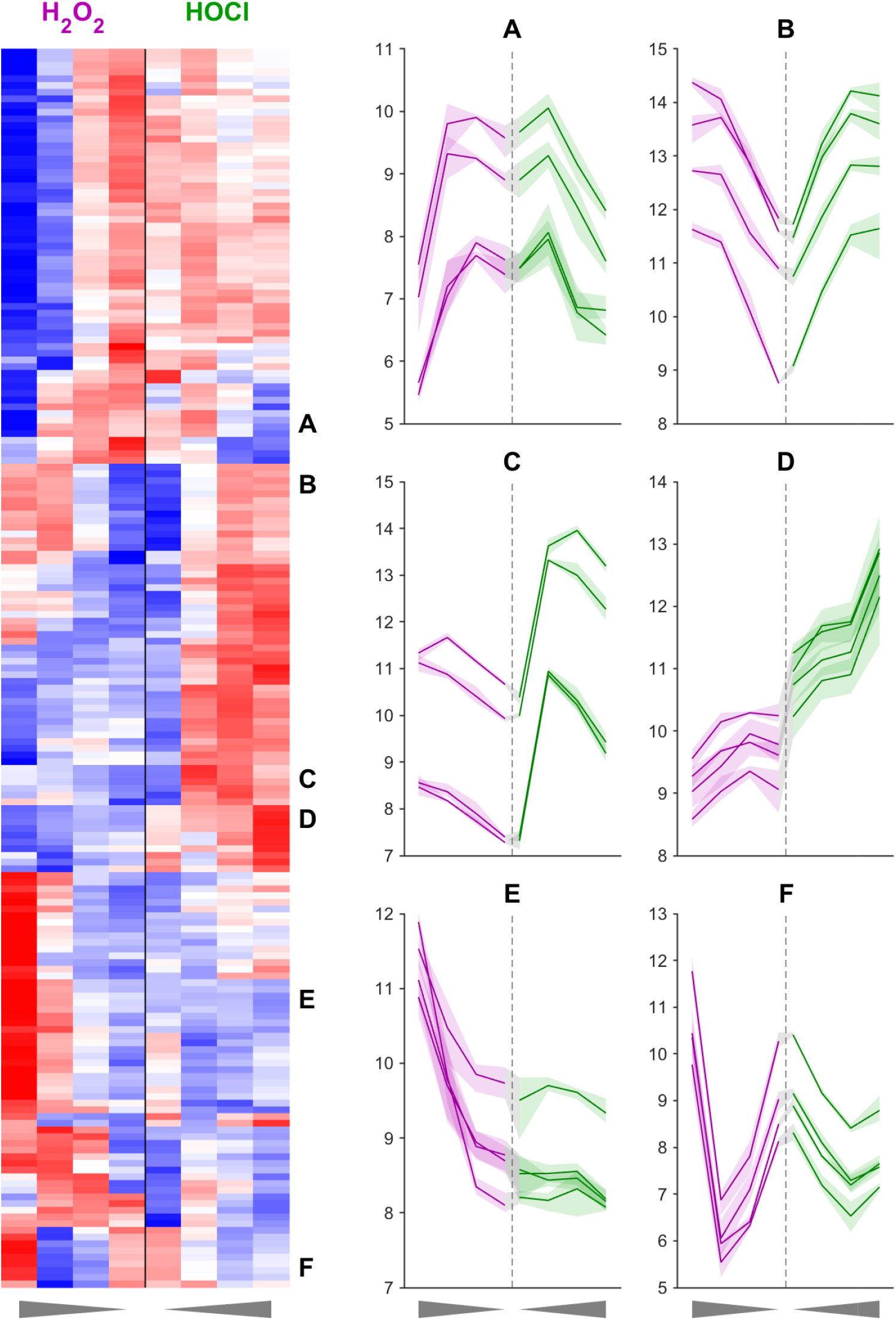
Clustering analysis of gene expression levels (FPKM) in anaerobically-grown *L. reuteri* ATCC PTA 6475 treated with 0.12 mM H_2_O_2_ (purple) or 1.25 mM HOCl (green) 0, 5, 15, and 30 minutes after stress treatment (times plotted outward from the center for each panel). Heat map showing row z-scores of rlog-transformed data (red = +2, blue = −2). Rows are 185 high-variance genes (out of 2010; variance > 0.5, after averaging triplicates), columns are treatment/time-points (left four columns are H_2_O_2_, right four columns are HOCl; time emanates from center along x-axis as indicated by triangles below plot). Rows are hierarchically clustered using Euclidean distance of z- score data and average linkage. Letters A-F are callouts of six example sets of four genes each whose expression data are shown in the six plots on the right. Solid lines are mean expression values (triplicates) and shaded regions indicate maximum and minimum values of triplicates for the corresponding treatment/time-point. Called-out genes are A: LAR_RS04660 (*ribD*), LAR_RS04665 (*ribE*), LAR_RS04670 (*ribBA*), LAR_RS04675 (*ribH*); B: LAR_RS03125, LAR_RS02625 (*oxc*), LAR_RS09770, LAR_RS10190; C: LAR_RS02280 (*copA2*), LAR_RS02285 (*copA3*), LAR_RS02290 (*copA*), LAR_RS09945; D: LAR_RS01550 (*cgl*), LAR_RS01555 (*cyuA*), LAR_RS01560 (*cyuB*), LAR_RS01565 (*cyuC*); E: LAR_RS08400, LAR_RS07525, LAR_RS07530, LAR_RS07535 (*cwlA*); F: LAR_RS05395 (*moaD*), LAR_RS05400 (*moaE*), LAR_RS05455, LAR_RS05460.

*L. reuteri*’s response to H_2_O_2_ was generally consistent with what has been previously observed in other catalase-negative Gram-positive bacteria (21, 23, 24, 31, 66–68). Highly upregulated genes included alkylhydroperoxidase (*ahpCF*)(69), NADH oxidase (*noxE*)(70), methionine sulfoxide reductase (*msrB*)(46), DNA repair genes (*uvrABD*, *xthA, umuC*)(71), predicted metal transporters (*pcl1*, *pcl2*), and the peroxide-sensing transcription factor PerR (68). The response to HOCl was also, in broad strokes, similar to that of previously characterized bacteria (22), in that upregulated genes included those involved in proteostasis (*groSL, clpE, hsp20/lo18*), metal stress *(pcl1, pcl2, copAR*), thioredoxins (*trxABD*), and cysteine and methionine synthesis (*cysK, metE*). Genes upregulated by both stressors included not only *msrB*, *ahpCF, perR*, and the predicted iron transporters encoded by *pcl1* and *pcl2*, but a variety of predicted sugar and amino acid transporters and metabolic enzymes (*oxc*, encoding oxalyl-CoA decarboxylase (72), for example). These may represent responses to changes in the nutritional environment *L. reuteri* encounters in the inflamed gut (6, 8).

### Redox-regulated transcription factors in *L. reuteri*

While many bacterial transcription factors have been described that respond to H_2_O_2_ and / or HOCl, *L. reuteri* encodes only a few homologs of known H_2_O_2_-detecting transcription factors (*e.g.* PerR and VicK (24, 67)) and no close homologs of any of the known HOCl-detecting transcription factors (22, 53–55, 68, 73). This suggested that among the 104 predicted transcription factors encoded by the *L. reuteri* genome (Table S3), there are likely to be novel redox-sensing regulators. To begin to assess this possibility, we performed clustering analysis on the expression of genes encoding transcription factors under both stress conditions (Fig. 5), reasoning that many bacterial transcription factors are autoregulated, and that changes in expression of transcription factors are useful signposts for identifying regulatory stress-response networks (53, 65, 74). We found genes encoding predicted transcription factors whose expression was activated by both H_2_O_2_ and HOCl (*e.g. perR*, *spxA*, LAR_RS09770), repressed by both H_2_O_2_ and HOCl (*e.g. kdgR, fabT*), activated only by H_2_O_2_ (*e.g. lexA*, LAR_RS07525), activated only by HOCl (*e.g. ctsR, copR*), repressed only by H_2_O_2_ (*e.g. sigH*, *rex*), and repressed only by HOCl (*e.g. malR3*, LAR_RS02755), indicating the presence of a complex regulatory response to both oxidants. Some of these regulators have known functions, which give useful insights into the *in vivo* effects of H_2_O_2_ and HOCl on *L. reuteri*. For example, only HOCl activated expression of *ctsR*, a conserved regulator of protein quality control in Gram positive bacteria (75, 76), consistent with the known ability of HOCl to unfold and aggregate proteins (77, 78) and the activation of the heat shock response in many species of HOCl-stressed bacteria (22). On the other hand, only H_2_O_2_ activated expression of the DNA-damage responsive *lexA* regulator (71), consistent with the known ability of H_2_O_2_ to damage DNA (23), and suggesting that HOCl does not cause DNA damage at the concentration used in this experiment. However, most of the transcription factors in *L. reuteri* have no known function, and the expression patterns of many of these genes were affected by the redox stress treatments. For example, the only alternative sigma factor (79) encoded in the *L. reuteri* genome (*sigH*) was downregulated strongly by H_2_O_2_, but unaffected by HOCl. We do not currently know what genes these uncharacterized regulators regulate, what role(s) they may play in surviving redox stress, or what transcription factor(s) are responsible for HOCl-specific regulation in *L. reuteri*.

**FIG 5.**
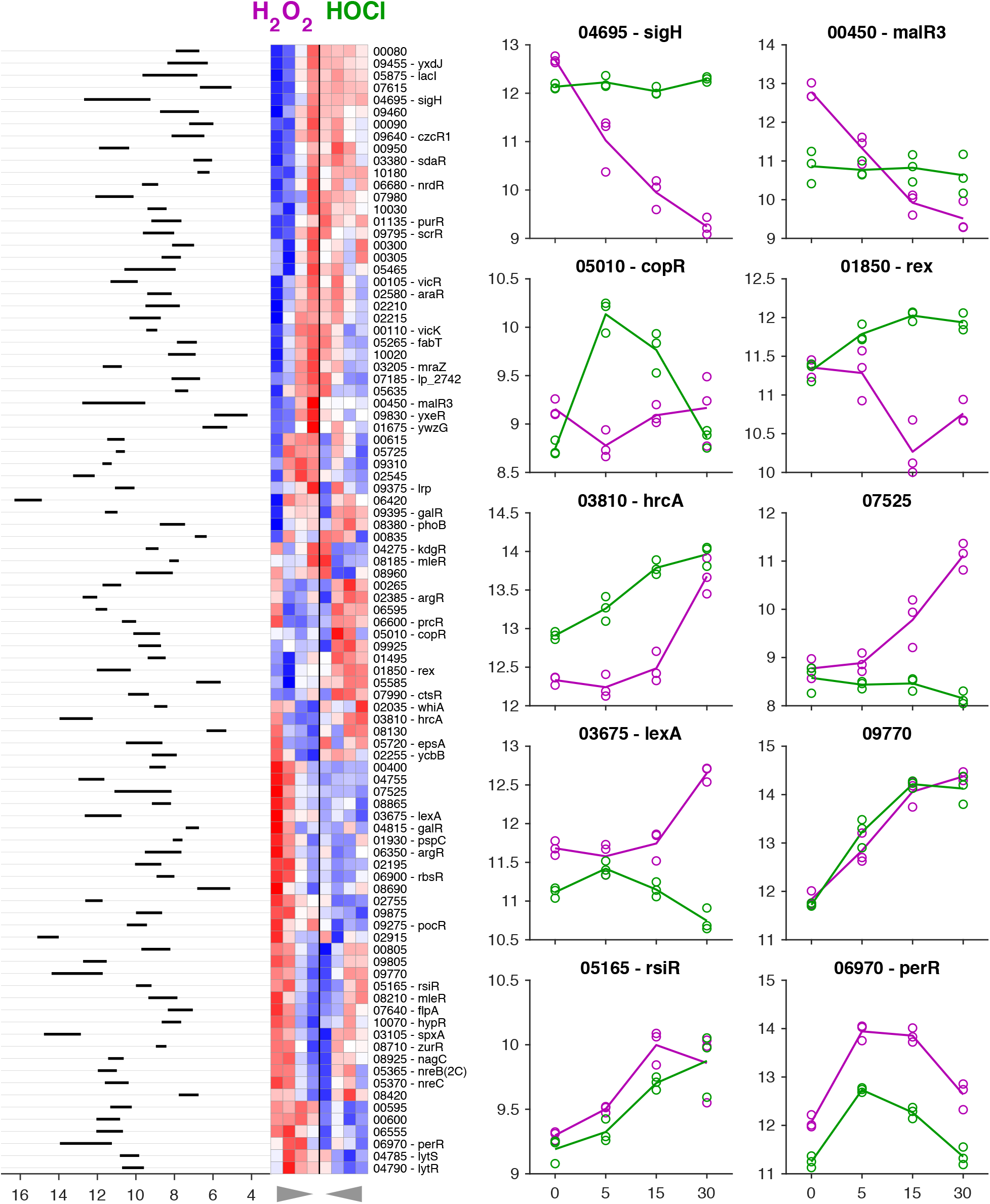
Clustering analysis of gene expression for transcription factors in anaerobically-grown *L. reuteri* ATCC PTA 6475 treated with 0.12 mM H_2_O_2_ (purple) or 1.25 mM HOCl (green) 0, 5, 15, and 30 minutes after stress treatment. Heat map showing z-scores of rlog-transformed data (red = +2, blue = −2). Rows are 93 transcription factors where the adjusted p-value was < 0.01 for any of the within-treatment baseline time comparisons, columns are treatment/time-points (left four columns are H_2_O_2_, right four columns are HOCl; time emanates from center along x-axis as indicated by triangles below plot). Rows are hierarchically clustered using Euclidean distance of z-score data and average linkage. rlog-transformed expression data are shown for ten example transcription factors to the right of the heat map (lines are means, circles are individual replicates). Expression range of averaged replicates shown to left of heat map. See Table S3 for expression and log_2_-fold change values for all predicted transcription factors.

### Oxygen affects H_2_O_2_- and HOCl-dependent gene expression in *L. reuteri*

We used quantitative RT-PCR to measure the changes in expression of selected genes in anaerobically grown *L. reuteri* treated with different concentrations of H_2_O_2_ and HOCl, at, above, and below the sublethal concentrations used in previous experiments (Fig. 6, left hand columns). We examined two genes strongly activated by both H_2_O_2_ and HOCl (*ahpF* and *pcl1*) and a gene repressed by both oxidants (*moeB*, encoding a subunit of molybdopterin synthase (80)) (Table S1) and confirmed the expected expression patterns. Interestingly, the genes differed in their dose-response patterns, with *moeB* roughly equally repressed at all H_2_O_2_ and HOCl concentrations, *ahpF* equally activated by all three H_2_O_2_ concentrations but activated more strongly by increasing doses of HOCl, and *pcl1* activated more strongly at lower doses of H_2_O_2_ and at higher doses of HOCl. RT-PCR of the *perR* and *sigH* regulator genes also recapitulated the expression patterns seen in RNA-seq (Fig. 5), although at higher HOCl concentrations (2.5 mM), *sigH* expression began to be repressed, indicating that its control is not strictly H_2_O_2_- specific. Finally, we examined expression of *rsiR*, a known regulator of anti-inflammatory mechanisms in *L. reuteri* (34, 35) that was modestly upregulated by both H_2_O_2_ and HOCl in the RNA-seq experiment (Fig. S4), but we did not observe activation of *rsiR* expression in this follow-up experiment, indicating that expression of *rsiR* may not genuinely be redox-regulated under anaerobic conditions.

**FIG 6.**
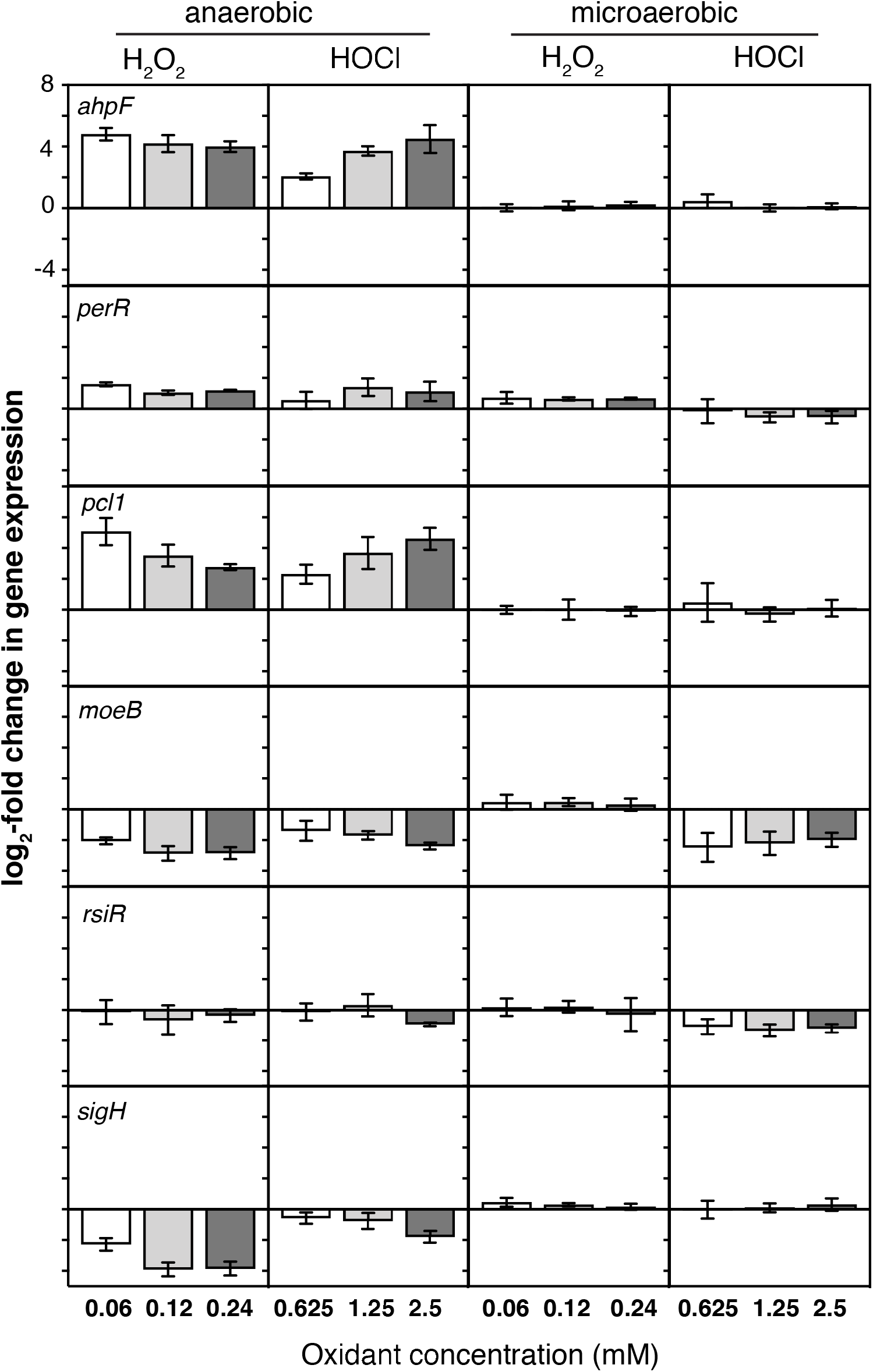
Dose-responsive control of gene expression by oxidative stress. *L. reuteri* ATCC PTA 6475 was grown anaerobically or microaerobically at 37°C to A_600_=0.3–0.4 in MEI-C and then treated with the indicated concentrations of H_2_O_2_ or HOCl. Change in expression of the indicated genes relative to untreated control cells was measured by quantitative RT-PCR (n=3, ±SD).

While the intestine is primarily an anaerobic environment (5), recent evidence suggests that inflammation, antibiotic treatment, and infection with enteric pathogens may increase the amount of oxygen available to microbes in the gut (6, 7). We therefore wanted to assess how much of an effect oxygen has on expression of redox regulated genes in *L. reuteri*. We repeated our RT-PCR experiment with microaerobic cultures, which were prepared aerobically and grown in full screw-cap tubes without shaking, low-oxygen conditions under which *L. reuteri* grows at a similar rate as under anaerobic conditions (data not shown). The results of this experiment (Fig. 6, right hand columns) revealed that the presence of even the low levels of oxygen expected in these cultures had large effects on redox-responsive gene expression. In contrast to what we observed anaerobically, expression of *ahpF, pcl1, moeB*, and *sigH* was unaffected by H_2_O_2_ under these conditions, and activation of *perR* was substantially reduced. HOCl activation of *ahpF, pcl1*, and *perR* expression was eliminated in the presence of oxygen, and both *moeB* and *rsiR* expression were HOCl-repressed. These results showed that oxygen can dramatically affect how bacteria regulate gene expression in response to inflammatory oxidants, and that studies of redox responses under aerobic growth conditions may not reflect how bacteria respond in anaerobic environments and *vice versa*.

### Identifying genes important for surviving oxidative stress in *L. reuteri*

Finally, we wanted to use the gene expression data generated above to begin identifying genes involved in protecting *L. reuteri* against the toxicity of H_2_O_2_ and HOCl, based on the simple hypothesis that genes strongly upregulated by a certain stress may be involved in protecting the cell against that stress (81). We were particularly interested in identifying genes encoding factors that protect *L. reuteri* against HOCl, since much less is known about HOCl defense in bacteria in general (22), and no previous studies have examined how lactic acid bacteria survive reactive chlorine stress. We therefore identified lethal doses of H_2_O_2_ and HOCl for *L. reuteri* (Fig. S3), and found that 1.5 mM H_2_O_2_ was sufficient to kill 99 – 99.9% of *L. reuteri* over the course of an hour both anaerobically and microaerobically. Anaerobic cultures of *L. reuteri* were killed by 7.5 mM HOCl, while microaerobic cultures died to a similar extent at only 2.5 mM HOCl, further demonstrating that oxygen influences how *L. reuteri* responds to toxic oxidants.

We constructed several strains containing null mutations of genes that we predicted to be involved in defense against either H_2_O_2_ or HOCl, based on known bacterial redox stress response mechanisms (22, 23, 53, 82) and on our transcriptomic data (Fig. S4, Table S4). We obtained mutants lacking *msrB*, *perR*, *sigH*, *rsiR*, *lo18* (*hsp20*), which encodes a small heat shock protein found only in lactic acid bacteria (59, 60) and whose expression was more strongly activated by HOCl than by H_2_O_2_, *ppk1* and *ppk2*, encoding two different kinases able to produce inorganic polyphosphate (polyP), which protects against HOCl-mediated protein damage in *E. coli* (57, 77, 83), *rclA*, encoding a conserved flavoprotein known to protect *E. coli* against HOCl (53), *hslO*, encoding the HOCl-activated chaperone Hsp33 (78), and LAR_RS09945, encoding a predicted oxidoreductase that was very strongly upregulated by HOCl, but not by H_2_O_2_. The ability of each of these strains to survive lethal oxidative stress was measured by comparison to the survival of the wild-type strain under the same conditions (Fig. 7).

**FIG 7.**
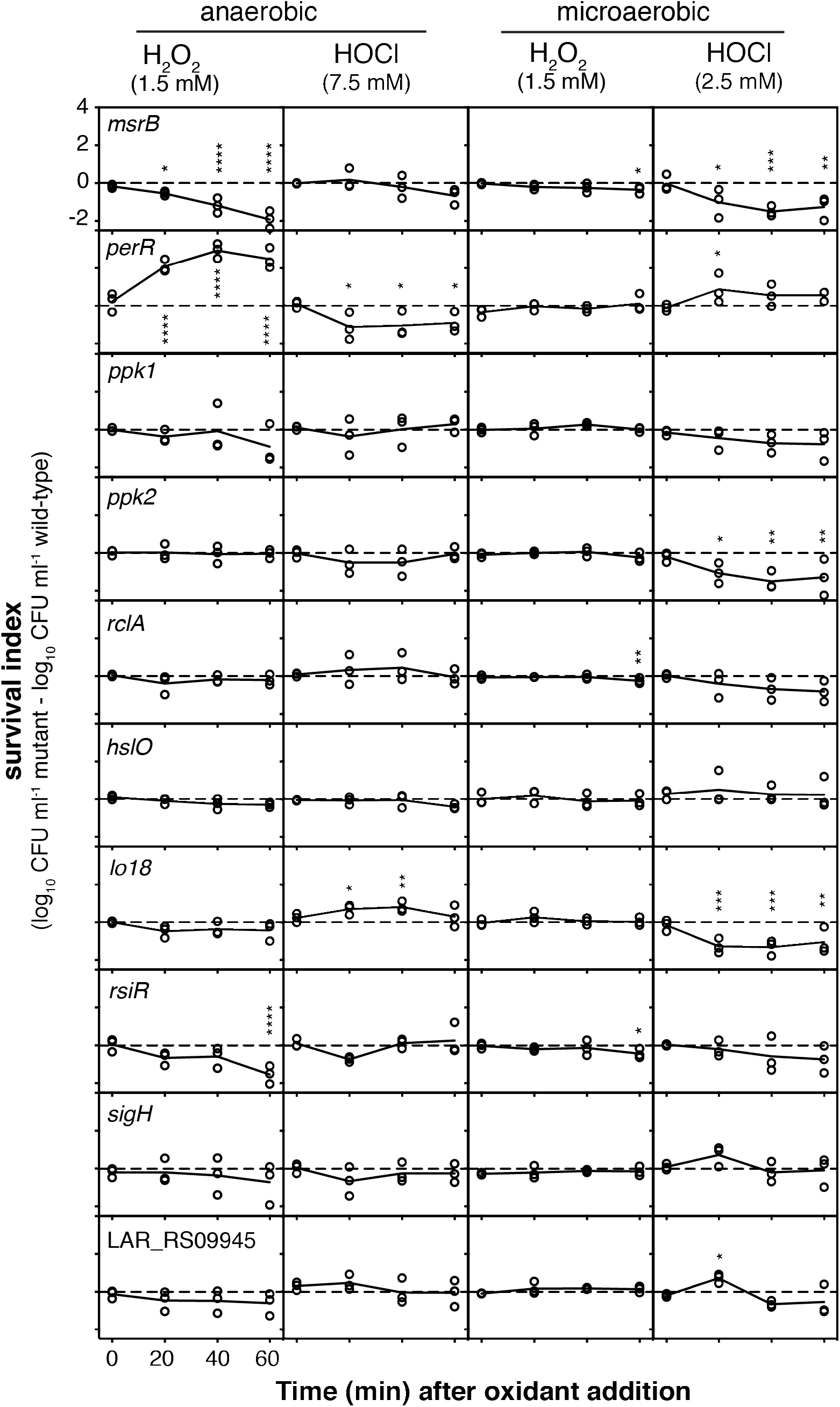
Genes regulated by inflammatory oxidants affect survival of lethal oxidative stress in *L. reuteri*. Survival index (log_10_ CFU ml^−1^ of mutant strain minus the log_10_ CFU ml^−1^ of wild-type strain from the same experimental replicate) of anaerobically or microaerobically-grown *msrB*, *perR*, *ppk1*, *ppk2*, *rclA*, *hslO*, *lo18*, *rsiR*, *sigH*, and LAR_RS09945 null mutants at the indicated time points after addition of 1.5 mM H_2_O_2_ or 7.5 or 2.5 mM HOCl. Asterisks indicate survival indices significantly different from zero at the indicated time point (two-way repeated measures ANOVA with Holm-Sidak’s multiple comparisons test, * = p<0.05, ** = p<0.01, *** = p<0.001, **** = p<0.0001).

Anaerobically, the *msrB* mutant was extremely sensitive to lethal H_2_O_2_ treatment, as expected (23, 46), and the *perR* mutant, which is expected to have constitutively high expression of peroxide defense genes (84), was significantly protected. A mutant lacking *rsiR* was significantly more sensitive to killing by H_2_O_2_, suggesting that despite the fact that its expression may not be controlled by this oxidant (Fig. 6), it is important for surviving H_2_O_2_ treatment (35). Surprisingly, only the *perR* mutant was significantly more sensitive than wild-type to killing by HOCl under anaerobic conditions. However, knocking out *lo18* had a significant and unexpected protective effect. This was particularly surprising since *lo18* expression was strongly upregulated in response to HOCl (Table S4). Under microaerobic conditions, the results of survival assays were considerably different. There were only minor differences in survival of a lethal dose of H_2_O_2_ in microaerobic cultures for any of the mutants, with *msrB*, *rsiR*, and *rclA* mutants showing very small but statistically significant defects in survival at the 1-hour time point. In contrast, there were much more substantial differences in survival of lethal HOCl stress under microaerobic conditions. The *msrB*, *perR, lo18* and *ppk2* mutants had significant defects in HOCl stress survival under these conditions, and the *rsiR, ppk1*, and *rclA* mutants were, on average, slightly more sensitive than the wild-type. The *perR* and LAR_RS09945 mutants were significantly protected at the 20 minute timepoint, but this effect was lost at later time points. There was no difference in HOCl survival between the wild-type and *sigH* or *hslO* mutants. These results further emphasize that oxygen concentration has dramatic effects on oxidative stress survival, and that it will therefore be important to quantify what oxygen levels gut bacteria are exposed to in inflamed and non-inflamed gut environments (5–7) to understand what genes are likely to play roles in ROS and RCS resistance *in vivo*.

Screening mutants lacking HOCl-induced genes has successfully identified HOCl resistance factors in other bacterial species (53, 54, 56, 65), but this strategy had limited success in *L. reuteri*. Neither *sigH* nor LAR_RS09945 mutations, for example, had any effect on survival of the stresses which regulate their expression. In future work, a genome-wide mutant screening approach (*e.g.* transposon sequencing)(52) may be valuable for identifying additional genes required for H_2_O_2_ and HOCl stress survival. Nevertheless, our targeted mutagenesis approach did allow us to identify several important players in oxidative stress resistance. Clearly, methionine sulfoxide reductase is a major contributor to the ability of *L. reuteri* to survive oxidative stress both anaerobically and microaerobically, consistent with its enzymatic activity (46) and known role in colonization (47, 48). While PerR is relatively unimportant microaerobically, anaerobically it plays an key role in regulating H_2_O_2_ resistance, as expected (68), although for unknown reasons it appears that the constitutive overexpression of H_2_O_2_-resistance genes expected in a *perR* mutant is detrimental in the presence of HOCl.

### *L. reuteri*-specific defenses against H_2_O_2_ and HOCl stress

The H_2_O_2_-sensitivity of the *rsiR* mutant was somewhat surprising, since this *L. reuteri*-specific gene has largely been characterized for its role in regulating the expression of the histamine-producing histidine decarboxylase locus of *L. reuteri*, where *rsiR* is essential for histamine-dependent anti-inflammatory phenotypes (34, 35). However, RsiR is a global regulator, activating and repressing transcription of 195 and 143 genes, respectively, many of which are involved in redox homeostasis (including *ahpC*, *perR*, and genes involved in cysteine and methionine synthesis)(35). It is currently unclear what signal(s) RsiR responds to, which RsiR-regulated genes contribute to H_2_O_2_ sensitivity, or what role H_2_O_2_ resistance plays in RsiR-dependent anti-inflammatory effects *in vivo*, and these are exciting questions for future research exploring the connections between inflammatory oxidants and anti-inflammatory probiotic mechanisms.

The small heat shock protein Hsp33 and the flavoprotein RclA are RCS-specific defense factors in *E. coli* (53, 78), so we were also surprised to find that mutations of these genes had no apparent effect on HOCl survival in *L. reuteri*, despite the fact that *rclA* expression was induced more strongly by HOCl treatment than by H_2_O_2_ (Table S4). This could be due to the redundant nature of RCS resistance mechanisms (22), or could reflect fundamental differences in RCS response between *L. reuteri* and *E. coli*. Supporting the second hypothesis is the fact that mutations in *lo18* and *ppk2*, genes not found in *E. coli*, had very strong effects on HOCl survival. Lo18 is a chaperone found only in a subset of *Lactobacillus* and *Oenococcus* species that stabilizes proteins and membranes under heat and ethanol stress conditions (59, 60). While this could easily explain how Lo18 protects *L. reuteri* against the protein-unfolding activity of HOCl, as we saw under microaerobic conditions, it is much less intuitive why the presence of Lo18 sensitizes *L. reuteri* to HOCl anaerobically, and more work will be needed to understand the mechanism underlying this effect. PolyP plays a role in stress resistance and probiotic phenotypes in several different *Lactobacillus* species (85–90). In *E. coli*, the polyP kinase PPK (homologous to *L. reuteri* PPK1) is required for HOCl resistance (77), but deletion of *ppk1* had only a modest, non-statistically significant effect on HOCl resistance in *L. reuteri*. In contrast, deletion of *ppk2*, which encodes an unrelated polyP kinase (PPK2) whose primary physiological role is generally thought to be in generating NTPs from NDPs or NMPs and polyP (57, 58), led to a highly significant defect in HOCl survival, albeit only in the presence of oxygen. Whether polyP production in response to HOCl stress is driven by PPK1 or PPK2 in *L. reuteri* remains to be determined, as does the relative importance of PPK2’s polyP- and NTP-synthesizing activities. PPK2 is not present in enterobacteria, but is found in many species of commensal bacteria (including lactobacilli, Bacteroidetes, and Clostridiacea)(58, 91, 92)

Our results clearly demonstrate that HOCl resistance in *L. reuteri* depends on different factors than in *E. coli* or *B. subtilis*. These differences may represent targets for differentially sensitizing gut bacteria to oxidative stress. Interestingly, the frontline IBD drug mesalamine has recently been shown to be an inhibitor of PPK1 (93), and it is tempting to speculate that mesalamine may therefore have a larger impact on the ability of enterobacteria to survive in the inflamed gut than on PPK2-encoding commensals, although more data will be needed to test this hypothesis.

## CONCLUSIONS

Manipulating the microbiome is likely to be a key element in future treatments for inflammatory diseases of the gut. Development of such treatments will require a sophisticated understanding of how gut bacteria respond to changes in their environment. The differences we have now begun to uncover in oxidative stress response between anti-inflammatory, health-associated bacteria and pro-inflammatory, disease-associated species may present opportunities for new therapies. We hope our results will ultimately make it possible to sensitize enterobacteria to inflammatory oxidants while simultaneously protecting the healthy gut community.

## MATERIALS AND METHODS

### Bacterial strains and growth conditions

All strains and plasmids used in this study are listed in Table 1. All *L. reuteri* strains were derivatives of strain ATCC PTA 6475 (Biogaia)(94). Strain 6475*rsiR*-Stop (35) was a gift from James Versalovic (Baylor College of Medicine), and plasmid pJP042 (*recT*^+^ *erm*^+^)(94) was a gift from Jan-Peter van Pijkeren (University of Wisconsin – Madison). *L. reuteri* was grown at 37°C in MEI broth (86) without added cysteine (MEI-C) or on solid De Man, Rogosa, and Sharpe (MRS) agar (Difco). Anaerobic cultures were incubated in an anaerobic chamber (Coy Laboratory Products) in an atmosphere of 90% nitrogen, 5% CO_2_, and 5% H_2_ or in Hungate tubes prepared, inoculated, and sealed in that chamber. Liquid media were made anaerobic before use by equilibration for at least 24 hours in the anaerobic chamber. MRS plates for colony forming unit (CFU) plate counts were incubated in sealed containers made anaerobic using GasPak^TM^ EZ sachets (Becton Dickinson). Microaerobic cultures were incubated aerobically without shaking in 16 x 125 mm screw-cap test tubes containing 15 ml of MEI-C. Details of H_2_O_2_ and HOCl stress treatments, transcript quantification, and phenotype analysis are described in the Supplemental Material.

**TABLE 1.** Strains and plasmids used in this study. Unless otherwise indicated, all strains were generated in the course of this work.

| Strain | Relevant Genotype | Source |
| --- | --- | --- |
| <u>*L. reuteri* strains:</u> |  |  |
| ATCC PTA 6475 | wild-type, human breast milk isolate | Biogaia,<br>(94) |
| 6475*rsiR*-Stop | *rsiR* (LAR_RS05165)<sup>G6C, G7C, T10A, A11T, C12G, A13T</sup> | (35) |
| MJG0562 | *ppk1* (LAR_RS01770)<sup>G94T, A95C, G96C, G97T, C98A</sup> |  |
| MJG0569 | *ppk2* (LAR_RS00075)<sup>A52C, C53T, G54T, G55T, C56A</sup> |  |
| MJG0570 | *rclA* (LAR_RS00915)<sup>C178A, A179G, T180C, G181T, G182A</sup> |  |
| MJG0977 | *msrB* (LAR_RS00975)<sup>G58T, T59C, T60C, A61T, C62A</sup> |  |
| MJG0979 | *hsIO* (LAR_RS01385)<sup>G106A, A107T, T108G, A109T, C110A</sup> |  |
| MJG1017 | *lo18* (LAR_RS07000)<sup>T43A, T44C, G45T, A46T, T47A</sup> |  |
| MJG1056 | LAR_RS09945<sup>C103A, C104T, A105G, G106T, A107G</sup> |  |
| MJG1278 | *sigH* (LAR_RS04695)<sup>G101A, G102A, C103T, C104A, G105A</sup> |  |
| MJG1573 | *perR* (LAR_RS06970)<sup>G10T, C11T, A12C, G13T, A14G</sup> |  |

### Molecular methods

Oligo-directed recombineering was used to construct null mutations in the chromosome of *L. reuteri* using the pJP042-encoded RecT recombinase as previously described (94). Null mutations were designed to incorporate in-frame stop codons near the 5’ end of each gene. Mutagenic primers used are listed in Table S5. Primers used for quantitative RT-PCR were designed with Primer Quest (www.idtdna.com; parameter set “qPCR 2 primers intercalating dyes” for qRT-PCR primer design) and are listed in Table S6. Additional primers for PCR amplification, screening, and sequencing were designed using WebPrimer (www.candidagenome.org/cgi-bin/compute/web-primer). All chromosomal mutations were confirmed by Sanger sequencing (UAB Heflin Center for Genomic Sciences).

### Data availability

All strains generated in the course of this work are available from the authors upon request. RNA sequencing data have been deposited in NCBI’s Gene Expression Omnibus (95) and are accessible through GEO Series accession number GSE127961 (https://www.ncbi.nlm.nih.gov/geo/query/acc.cgi?acc=GSE127961).

## ACKNOWLEDGEMENTS

We thank Drs. Rob Britton and James Versalovic (Baylor College of Medicine) and Dr. J.P. van Pijkeren (University of Wisconsin – Madison) for strains and plasmids. This project was supported by University of Alabama at Birmingham Department of Microbiology startup funds and NIH grant R35GM124590 (to M.J.G). The authors have no conflicts of interest to declare.

